# Cohesin-dependent and independent mechanisms support chromosomal contacts between promoters and enhancers

**DOI:** 10.1101/2020.02.10.941989

**Authors:** Michiel J. Thiecke, Gordana Wutz, Matthias Muhar, Wen Tang, Stephen Bevan, Valeriya Malysheva, Roman Stocsits, Tobias Neumann, Johannes Zuber, Peter Fraser, Stefan Schoenfelder, Jan-Michael Peters, Mikhail Spivakov

**Affiliations:** Nuclear Dynamics Programme, Babraham Institute, Cambridge CB22 3AT, UK; Research Institute of Molecular Pathology, Vienna Biocenter, Vienna, Austria.; MRC London Institute of Medical Sciences, London W12 0NN, UK; Institute of Clinical Sciences, Faculty of Medicine, Imperial College, London W12 0NN, UK; Department of Biological Science, Florida State University, Tallahassee, FL 32301; Epigenetics Programme, Babraham Institute, Cambridge CB22 3AT, UK

**Keywords:** Cohesin, CTCF, promoter capture Hi-C, SLAM-seq, promoter-enhancer interactions, transcriptional regulation

## Abstract

It is currently assumed that 3D chromosomal organisation plays a central role in transcriptional control. However, recent evidence shows that steady-state transcription of only a minority of genes is affected by depletion of architectural proteins such as cohesin and CTCF. Here, we have used Capture Hi-C to interrogate the dynamics of chromosomal contacts of all human gene promoters upon rapid architectural protein degradation. We show that promoter contacts lost in these conditions tend to be long-range, with at least one interaction partner localising in the vicinity of topologically associated domain (TAD) boundaries. In contrast, many shorter-range chromosomal contacts, particularly those that connect active promoters with each other and with active enhancers remain unaffected by cohesin and CTCF depletion. We demonstrate that the effects of cohesin depletion on nascent transcription can be explained by changes in the connectivity of their enhancers. Jointly, these results provide a mechanistic explanation to the limited, but consistent effects of cohesin and CTCF on steady-state transcription and point towards the existence of alternative enhancer-promoter pairing mechanisms that are independent of these proteins.

## Introduction

DNA regulatory elements such as enhancers play key roles in transcriptional control, particularly for developmental and stimulus-response genes (Long et al., 2016; Shlyueva et al., 2014). Many enhancers are located large distances (up to megabases) away from their target promoters, and are brought into physical proximity with them through DNA looping interactions (Göndör and Ohlsson, 2018). 3D chromosomal architecture and its regulators are therefore thought to be important for transcriptional control.

The cohesin complex and regulatory proteins that control cohesin-DNA interactions are critical for shaping chromosomal architecture (Merkenschlager and Nora, 2016). According to the current understanding, cohesin is involved in extrusion of DNA loops in interphase nuclei (Davidson et al., 2019; Kim et al., 2019), additionally to its well-characterised role in holding sister chromatids together in mitosis (Alipour and Marko, 2012; Fudenberg et al., 2016; Goloborodko et al., 2016; Sanborn et al., 2015). Extruding DNA loops are confined by chromatin boundary elements, in particular by the binding of zinc-finger protein CTCF to its two recognition motifs on the DNA in convergent orientation (Rao et al., 2014; Vietri Rudan et al., 2015). These loops underpin the formation of topologically associated domains (TADs (Dixon et al., 2012; Nora et al., 2012; Sexton et al., 2012)) and substructures such as insulated neighbourhoods (INs) (Hnisz et al., 2016; Nuebler et al., 2018; Szabo et al., 2019). Depletion of either cohesin leads to rapid dissolution of TADs, whereas inactivation of CTCF reduces the strength of TAD boundaries (Nora et al., 2017; Rao et al., 2017; Schwarzer et al., 2017; Wutz et al., 2017). In contrast, the higher-order levels of chromosomal organisation, such as A/B compartments that broadly separate active and inactive chromatin, remain largely unaffected by cohesin and CTCF depletion (Gassler et al., 2017; Nora et al., 2017; Rao et al., 2017; Schwarzer et al., 2017; Seitan et al., 2013; Sofueva et al., 2013; Vian et al., 2018; Wutz et al., 2017; Zuin et al., 2014).

Chromosomal domains such as TADs and INs constrain promoter-enhancer interactions (Bonev and Cavalli, 2016; Sun et al., 2019), albeit incompletely (Freire-Pritchett et al., 2017; Javierre et al., 2016), and perturbations of their boundaries can lead to gene misregulation in development and in cancer (Lupiáñez et al., 2016; Robson et al., 2019). However, genes sharing these domains may have different regulatory wiring and expression patterns (Hnisz et al., 2016; Schoenfelder and Fraser, 2019). In addition, active enhancers might themselves function as TAD boundary elements (Barrington et al., 2019; Bonev et al., 2017), and loop domains connecting superenhancers recover more rapidly upon cohesin depletion (Rao et al., 2017). These observations suggest that factors acting *in cis* to DNA regulatory elements are involved in establishing their chromosomal interactions. The interplay between higher-order domains and specific regulatory chromosomal interactions such as those between enhancers and their target promoters is not fully understood.

Consistent with a role for cohesin in shaping gene regulatory architecture, its depletion prevents adequate activation of inducible genes (Cuartero et al., 2018). Surprisingly however, steady-state gene expression levels are less affected upon architectural protein depletion (Busslinger et al., 2017; Haarhuis et al., 2017; Nora et al., 2017; Rao et al., 2017; Schwarzer et al., 2017; Seitan et al., 2013; Sofueva et al., 2013). This suggests that gene expression may be maintained by mechanisms independent of these architectural proteins, but whether this involves continued input from enhancers remains unclear.

Foundational studies of the effects of architectural protein depletion on 3D chromosomal architecture were performed using Hi-C (Lieberman-Aiden et al., 2009). While this is a powerful method for global detection of chromosomal conformation, the complexity of Hi-C sequencing libraries limits the coverage and resolution of data obtained using this technology, making the robust analysis of specific enhancer-promoter interactions challenging. Combining Hi-C with sequence capture (Capture Hi-C) makes it possible to mitigate this limitation by selectively enriching Hi-C libraries for interactions involving, at least on one end, regions of interest such as gene promoters (Mifsud et al., 2015; Sahlén et al., 2015; Schoenfelder et al., 2015a). The fact that this approach does not depend on proteins bound to either interaction partner makes it particularly suitable for studying interactions where these proteins are either unknown or ectopically depleted.

Here, we use Capture Hi-C to study the effects of architectural protein depletion on promoter interactions. We show that while a majority of promoter interactions dissolve upon cohesin and CTCF depletion, large numbers of interactions remain unaffected, and some interactions are gained in these conditions. Interactions that are lost, gained and maintained upon cohesin depletion have distinct properties with respect to localisation within TADs, interaction distance and/or the identity of associated proteins. We further demonstrate that the effects of cohesin depletion on nascent transcription of specific genes (as measured by SLAM-seq) can be generally explained by changes in the connectivity of their active enhancers. These results provide a mechanistic explanation to the limited, but significant effects of cohesin and CTCF perturbations on gene expression and suggest the existence of alternative mechanisms supporting promoter-enhancer interactions.

## Results

### Extensive rewiring of promoter interactions upon rapid depletion of architectural proteins

To study the effects of architectural protein depletion on promoter interactions, we took advantage of HeLa cells, in which all alleles of either cohesin subunit *SCC1* or *CTCF* were tagged with a minimised auxin-inducible degron (Morawska and Ulrich, 2013) and an mEGFP reporter (SCC1-AID and CTCF-AID cells, respectively). Additionally, these cells stably express *Oryza sativa* Tir1 protein required for proteasome targeting of AID-tagged proteins (Nishimura et al., 2009). We previously showed that SCC1 and CTCF are rapidly degraded in these cell lines within 20 min of auxin treatment (Wutz et al., 2017).

We performed high-resolution Promoter Capture Hi-C (PCHi-C) in cell-cycle synchronised, auxin-treated SCC1-AID and CTCF-AID cells and untreated controls. In addition, to compare the effects of depletion of these proteins with cell-cycle effects, we performed PCHi-C in intact HeLa cells synchronised in G1, G2 and mitosis. Finally, we profiled cells, in which cohesin release factor WAPL was depleted with RNAi (Tedeschi et al., 2013; Wutz et al., 2017). Two biological replicates of each condition were sequenced, aligned and filtered using the HiCUP pipeline (Wingett et al., 2015) to a median coverage of ∼95 M valid read pairs each, with the overall coverage of ∼1.5 billion valid read pairs. Given the ∼17 fold enrichment for captured promoter interactions compared with conventional Hi-C, the combined coverage of promoter interactions in our dataset is equivalent to that achievable with ∼25 billion Hi-C read pairs. Using the CHiCAGO pipeline (Cairns et al., 2016), we detected on average ∼100,000 significant interactions between promoters and promoter interacting regions (PIRs) in unperturbed cells (see Figs 1A and 6A for examples). The numbers of significant interactions were however markedly lower in architectural protein-depleted cells compared with controls, with a particularly strong effect for cohesin (118,074 SCC1 control and 61,702 SCC1 depleted, respectively; Fig. S1).

Clustering of interactions based on CHiCAGO scores identified 13 coherent clusters denoted A to M (Fig. 1B; see Methods for details). Clusters B, C, F were characterised by loss of interaction signal upon cohesin and CTCF depletion, while interactions in cluster D were sensitive to degradation of cohesin, but not CTCF. Notably, all of these four clusters also showed loss of promoter interaction signal in mitosis, in which the majority of these proteins are thought to be released from chromosomes. In addition, promoter interactions in these clusters appeared generally stronger in unmodified HeLa cells compared with AID-modified, but auxin-untreated cells (“controls”, Fig. 1B). This is likely due to the baseline residual activity of the AID system without auxin observed previously (Wutz et al., 2017).

**Figure 1.**
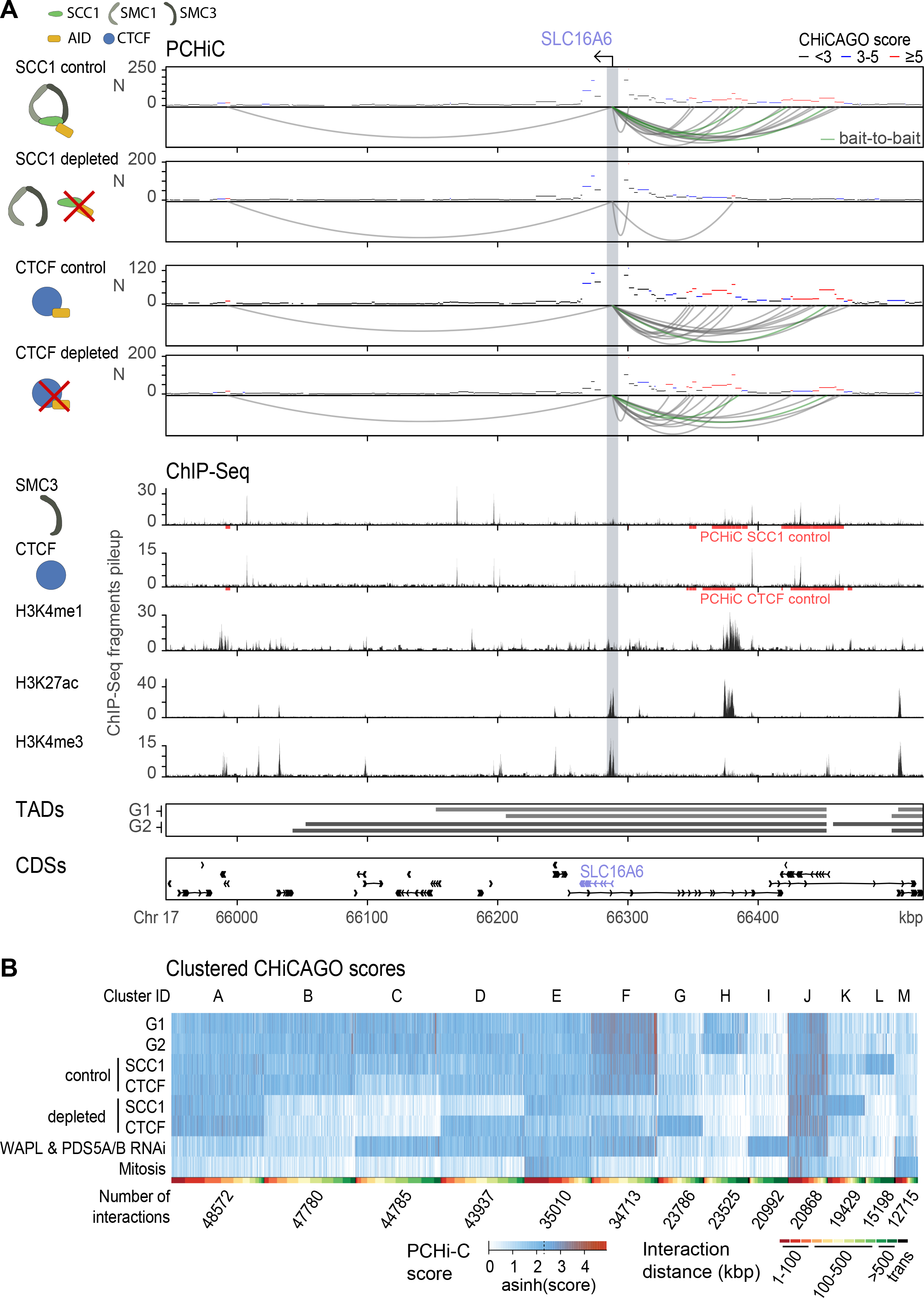
Promoter interaction rewiring upon architectural protein depletion. (A) Chromosomal interactions detected for *SLC16A6* promoter by PCHi-C. The top four tracks show PCHi-C interaction profiles with interaction arcs for the conditions: SCC1 control, SCC1 depleted, CTCF control, CTCF depleted. The following five tracks show ChIP-seq pileups in interphase for the targets: SMC3, CTCF, H3K4me1, H3K27ac, and H3K4me3. The bottom two tracks show TAD intervals in cell cycle stages G1 and G2, and coding sequences (GRCh37). Upon SCC1 depletion, the promoter of *SLC16A6* loses the majority of its promoter interactions. Upon CTCF depletion however, its promoter interactions appear to be preserved. The cohesin subunit SMC3 is present at many of the promoter interacting regions (PIRs), whereas H3K4me1 and H3K4me3 are present at a selection of PIRs. The promoter of *SLC16A6* and the majority of its interactions are located within the same TAD. (B) Clustered PCHiC interaction scores from the following conditions: synchronised interphase cell cycle stages G1 and G2, AID -Auxin control samples for SCC1 and CTCF, AID +Auxin depletions of SCC1 and CTCF, RNAi for the cohesin loading factor WAPL and its cofactors PDS5A and PDS5B, and finally synchronised cells in the mitotic prometaphase. A coherent partitioning with thirteen K-means clusters was found, which reveals promoter interactions that are dependent (clusters B, C, F) as well as independent (clusters A, E, J) from cohesin and CTCF. Interactions in cluster D are lost upon cohesin but not CTCF depletion.

Strikingly, promoter interactions in clusters A, E, J were maintained upon cohesin and CTCF depletion (Fig. 1B). Some of these interactions showed sensitivity to WAPL depletion and were lost in mitosis (cluster A), while others were generally retained in all analysed conditions (clusters E and J). Finally, distinct subsets of promoter interactions were gained depending on the depleted protein (SCC1, cluster K, CTCF, cluster G).

Jointly, these results point to both the shared and specific effects of cohesin and CTCF depletion on promoter wiring. In addition, they suggest that large numbers of promoter interactions are retained in these conditions, and these interactions appear generally stable throughout interphase.

Guided by the exploratory observations from cluster analysis, we next sought to define robust subsets of promoter interactions that are lost, maintained and gained upon cohesin and CTCF depletion with high confidence. For this, we additionally took advantage of our recently developed differential calling pipeline for PCHi-C data, Chicdiff (Cairns et al., 2019). Integrating Chicdiff and clustering results, we defined 36,174 lost; 12,978 maintained; and 2,484 gained interactions upon cohesin depletion (Figs 2A-C and S1C). The remaining promoter interactions were not assigned to any category at the predefined degree of confidence (see Methods for details). Notably, in an appreciable minority of cases, different interactions of the same regions (either promoters or PIRs) showed different dynamics upon cohesin depletion. For example, 1259/3412 (36.9%) of baited promoters whose interactions were lost upon cohesin depletion additionally had interactions that were either maintained or gained in cohesin-depleted cells (Fig. 2D). The same was true for 840/9324 (9%) of PIRs (Fig. 2E). Performing the same analysis for CTCF revealed considerably smaller numbers of significantly affected interactions, both overall (17,645 lost, 13,703 maintained and 1,663 gained, respectively) and on average per promoter (Fig. S1C).

**Figure 2.**
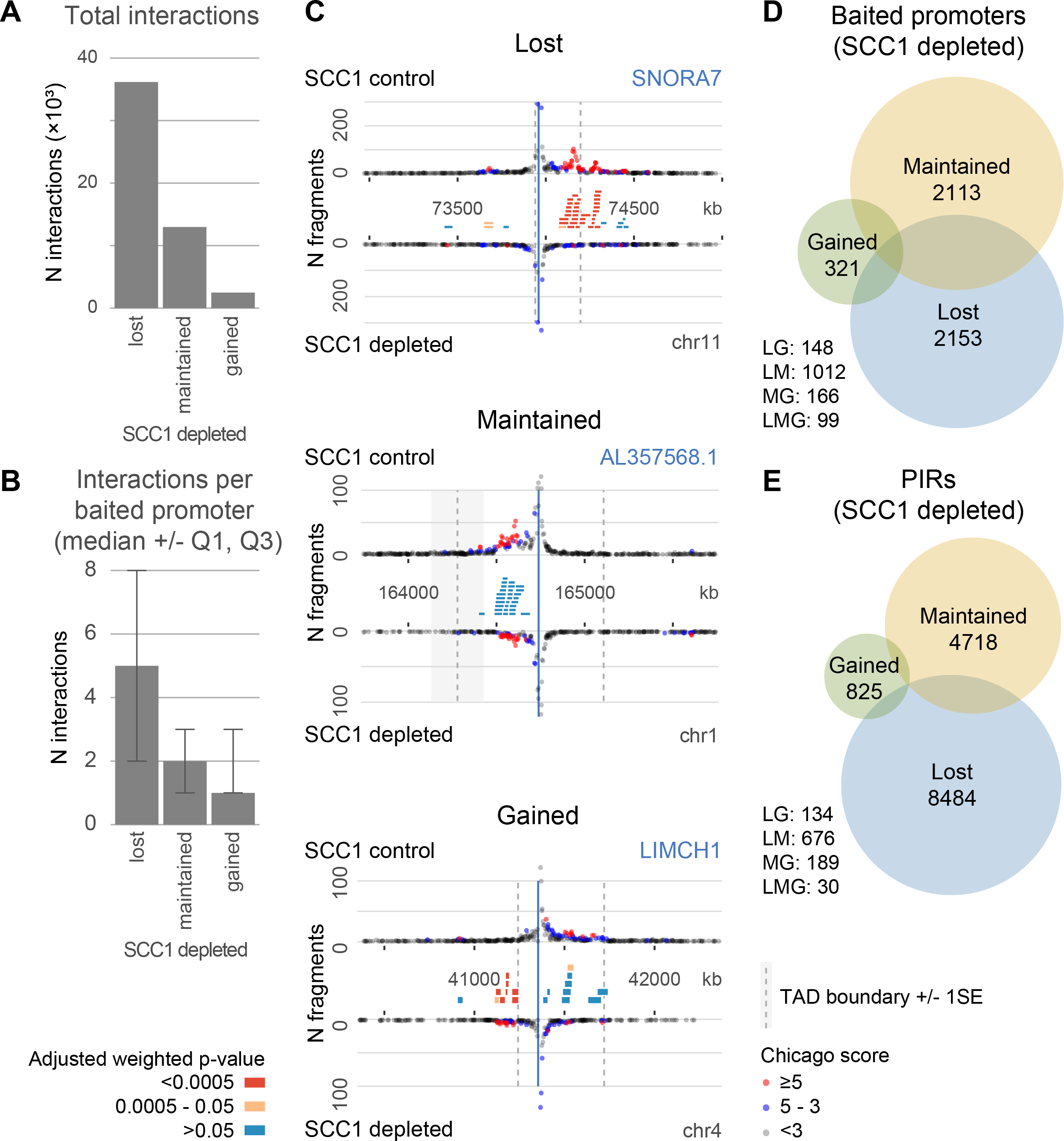
Lost, maintained and gained promoter interactions upon cohesin depletion. (A) Overview of the numbers of lost, maintained and gained promoter interactions. (B) Median number of lost, maintained and gained promoter interactions per baited promoter. Error bars indicate the interquartile range. (C) Illustrative examples of promoters with lost, maintained and gained interactions upon cohesin depletion. The blue bar indicates the baited restriction fragment containing a promoter. Red circles indicate significant promoter interactions. Red rectangles indicate significantly differential interactions between conditions (SCC1 control and depleted). The boundaries of the most proximal TAD are shown as dotted grey lines with a shaded area that represents +/-1SE over four interphase TAD calls. (D, E) Rewiring of interactions per baited promoter (D) and PIR (E).

The less pronounced effects of CTCF compared with cohesin depletion on promoter interactions are consistent with the hypothesis that cohesin is the primary factor in facilitating long-range interactions, while CTCF is often, but not always, required for cohesin positioning on the chromatin (Hansen et al., 2017; Nora et al., 2017; Rubio et al., 2008; Schmidt et al., 2010; Wutz et al., 2017). It cannot be ruled out however that the observed differences could also be due, at least in part, to the residual levels of CTCF previously detected upon auxin treatment in this system (Wutz et al., 2017). We therefore focused on cohesin depletion for the remainder of the study.

### TAD boundaries anchor cohesin-dependent and constrain cohesin-independent promoter interactions

Cohesin inactivation rapidly leads to a near-complete loss of TADs (Rao et al., 2017; Schwarzer et al., 2017; Wutz et al., 2017). We therefore asked how promoter interactions that were lost, maintained or gained upon cohesin depletion localised with respect to TAD boundaries. To address this, we focused on promoter interactions from either of these classes, for which at least one partner mapped within a TAD (representing 62.1% total). We partitioned the interactions into the following three categories based on the location of one interaction partner (i.e., either a baited promoter or PIR) relative to its nearest TAD boundary: boundary-proximal (mapping within 0-20% of the TAD length on either side), intermediate (20-40% TAD length), and mid-TAD (40-60% TAD length). For each class (lost, maintained, gained) and category (proximal, intermediate, mid-TAD), we then obtained the distribution of the locations of the second interaction partner relative to the boundaries of the respective TAD (Fig. 3A, summarised in Fig. 3B). We performed this analysis for interaction categories defined on the basis of either promoter or PIR locations, with similar results (Fig. 3AB).

**Figure 3.**
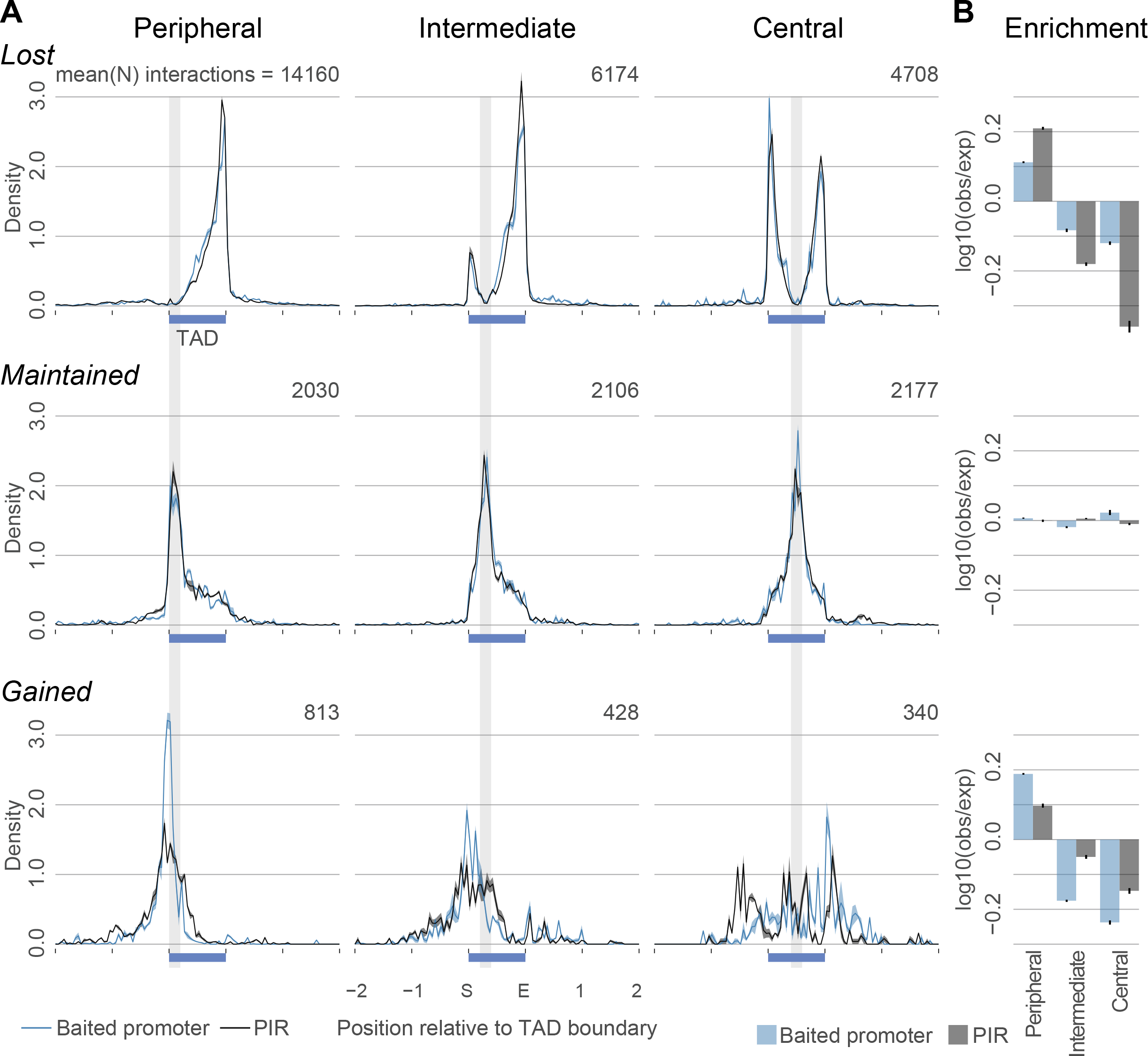
Localisation of promoter interactions that are lost, maintained and gained upon cohesin depletion with respect to TAD boundaries. (A) Promoter interaction frequency profiles. Promoter interactions that are lost, maintained or gained upon cohesin subunit SCC1 depletion are shown in the top, middle and bottom rows, respectively. Vertical grey bars represent the viewpoint window which encompasses TAD boundary proximal (left column), intermediate (middle column) and TAD central (middle column) positions. Horizontal blue bars represent TAD intervals. Black lines: distributions of locations of PIRs whose baited promoters map within the viewpoint window. Blue lines: distributions of locations of baited promoters whose PIRs map within the viewpoint window. The shaded area around the profile lines represents +/-1SE over four sets of interphase TAD calls. (B) Barcharts showing the enrichment of baited promoters and PIRs at specific windows within TADs.

As can be seen in Fig. 3 (top row), promoter interactions that were lost upon cohesin depletion tended to have at least one partner located in the vicinity of a TAD boundary, with the other interaction partner confined to the same TAD. Notably, for both the “boundary-proximal” and “intermediate” lost interactions, the second interaction partner most commonly localised near the “opposite”, distal boundary as opposed to the nearest one (Fig. 3A, top row – left and middle plots). In contrast, promoter interactions maintained upon cohesin depletion, while also constrained by TAD boundaries, showed no preference for the location of either partner within the same TAD (Fig. 3, middle row). Notably, unlike the lost promoter interactions, maintained interactions spanned relatively small proportions of the TADs’ lengths, irrespectively of their location relative to TAD boundaries (Fig. 3A, middle row). Finally, promoter interactions that were gained upon cohesin depletion were enriched around TAD boundaries and spanned relatively small proportions of the TADs’ lengths (Fig. 3AB, bottom row). The gained promoter interactions tended to cross the native TAD boundaries, indicating that they were likely enabled by the dissolution of TADs upon cohesin depletion. We have confirmed the TAD boundary constraint of lost and maintained interactions and the prevalence of TAD boundary-crossing gained interactions formally using logistic regression, accounting for interaction distance (Fig. S2). Jointly, these results suggest that cohesin-dependent (“lost”) promoter interactions are anchored to TAD boundaries, while cohesin-independent (“maintained”) promoter interactions are constrained by TADs, and those gained upon cohesin depletion are enabled by TAD dissolution.

### Cohesin-dependent and independent promoter interactions have distinct properties

We asked what features of promoter interactions associated with their dependence on or independence of cohesin. First, we found a striking association between longer interaction distances and cohesin dependence (Kruskal-Wallis test p<2.2e-16, Fig. 4A), which was observed even for the same PIRs involved in both lost and maintained interactions (Wilcoxon test p=2.2×10^-16^, Fig. 4B). We then assessed the binding of cohesin and CTCF to the interacting fragments (baits and PIRs), using previously published ChIP datasets for these (ENCODE Project Consortium, 2012; Wutz et al., 2017)(Wutz et al., 2017) Both baits and PIRs of lost interactions were selectively enriched for the binding of cohesin and CTCF compared with maintained and gained interactions (log odds ratio LOR = 1.052 and LOR = 1.360 at baits and LOR = 0.526 and LOR = 0.766 at PIRs respectively), demonstrating that architectural proteins likely mediate these interactions via their direct binding to the interaction partners in *cis* (Fig. 4C). We also assessed the binding of the active chromatin marks H3K4me1 (Kuznetsova et al., 2015; Nilson et al., 2017) and H3K4me3 (ENCODE Project Consortium, 2012; Liang et al., 2015) in the same way. Strikingly, we found that these marks showed the highest enrichment at the PIRs of maintained interactions and were relatively depleted at the lost interactions (Fig. 4C).

**Figure 4.**
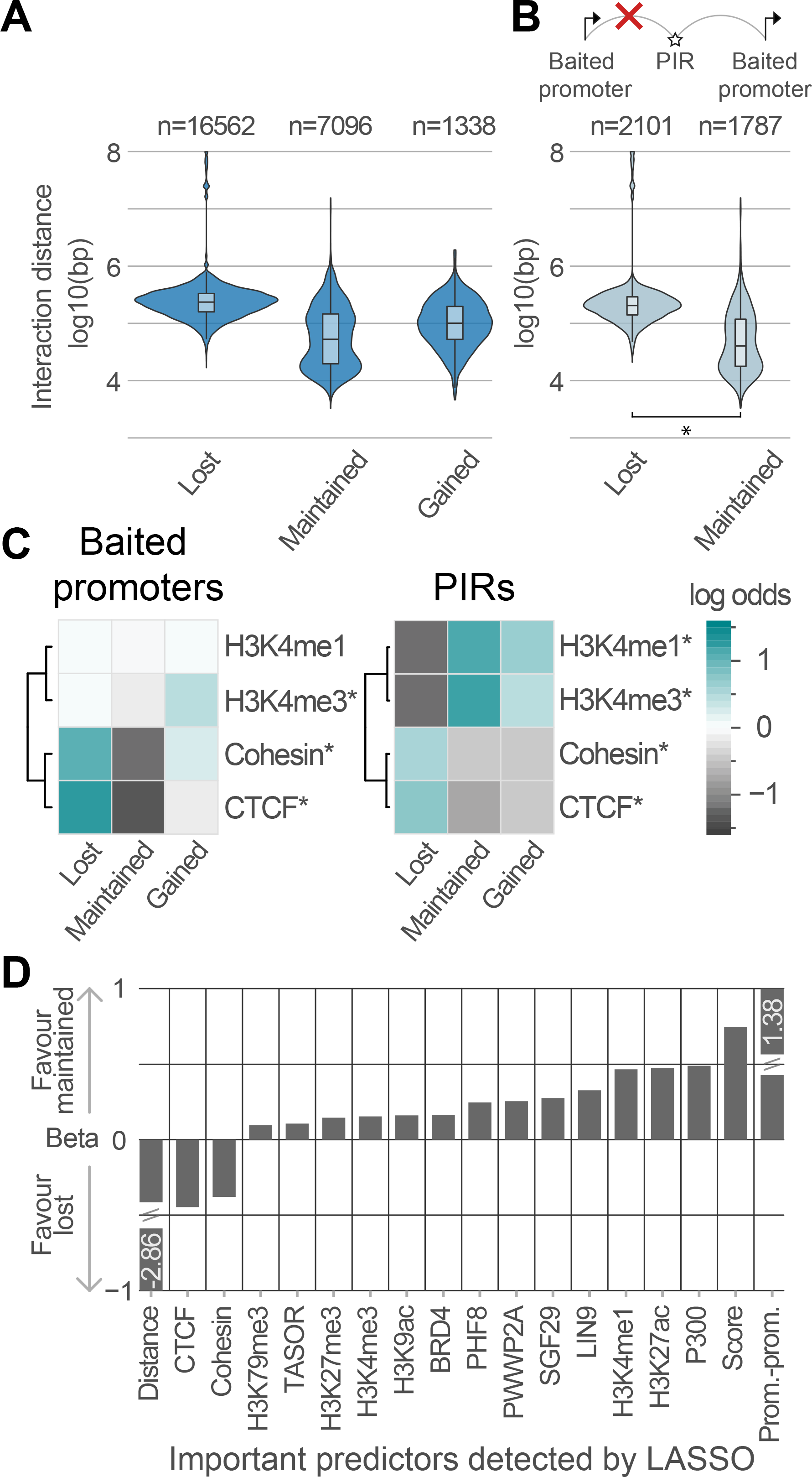
Chromatin features of cohesin-dependent and independent promoter interactions. (A) Violin plots showing the log_10_ genomic distance of lost, maintained and gained promoter interactions upon SCC1 depletion. (B) Violin plots showing the log_10_ genomic distance of lost (cohesin-dependent) and maintained (cohesin-independent) promoter interactions whose PIRs are engaged in the interactions of both types. * Wilcoxon test p=2.2×10^-16^. (C) Heatmaps showing log-odds ratios of restriction fragments (baited promoters, left; and PIRs, right, respectively) having strong ChIP-seq signals for active histone modifications and cohesin and CTCF binding. Signals marked with asterisks show significant differences across the lost, gained and maintained interaction categories (FDR-corrected Fisher test p-values < 0.05). (D) Barchart showing the regression coefficients (betas) of important predictors of maintained versus lost interactions identified by LASSO logistic regression. See Methods and Fig. S3 for details.

Next, focusing on the cohesin-dependent (“lost”) and independent (“maintained”) interactions, we expanded the panel of PIR-bound factors to ChIP datasets for 51 transcription factors (TFs) and 11 histone modifications in HeLa cells available from Cistrome and ENCODE projects (ENCODE Project Consortium, 2012; Zheng et al., 2019). We processed these data uniformly and devised a strategy to assess the signals at PIRs for each dataset in a consistent manner (see Methods for details). To delineate factors having the strongest association with cohesin dependence, we have employed LASSO logistic regression – a variable selection approach, in which explanatory variables with less important contributions to the outcome are removed from the model by shrinking their regression coefficients towards zero. The binary outcome variable reflected the maintenance versus loss of promoter interactions upon cohesin depletion. As explanatory variables, we used the full set of TFs and histone marks at PIRs. Additionally, we included interaction distance, interaction strength (expressed as CHiCAGO score), and whether an interaction connects two baited promoter regions.

The regression coefficients of the TFs and histones showing the strongest effects over a range of values of shrinkage parameter λ (from the most to the least restrictive) are shown in Fig. S3, and barplot in Fig. 4D summarises the effects of factors having significant associations with lost vs maintained interactions at an optimised level of shrinkage (see Methods for details). Interaction distance showed the strongest association with interactions lost upon cohesin depletion, followed by the binding of cohesin and CTCF (Fig. 4D). In turn, the PIRs of cohesin-independent (“maintained”) interactions showed a strong positive association with promoter-promoter contacts, stronger interaction strength, as well as features of active chromatin, including the histone marks H3K4me1, H3K4me3, H3K27ac and H3K9ac. These PIRs also preferentially recruited proteins associated with transcriptional activation, such as histone acetyltransferase p300, H3K9 demethylase PHF8, BET family protein BRD4, and LIN9 transcriptional regulator (Fig. 4C). We also found an association with H3K27me3 mark linked with Polycomb Repressor Complexes (Fig. 4C) that have known effects on chromosomal architecture (Cheutin and Cavalli, 2019).

We next asked whether cohesin-independent interactions were stabilised by transcription. To address this, we treated cells with RNA polymerase II (Pol II) inhibitor triptolide and compared their promoter interaction profiles with untreated cells. Triptolide treatment did not result in global changes in promoter interaction profiles. Furthermore, triptolide-induced changes in interaction scores were somewhat more pronounced for interactions also affected by cohesin depletion (Fig. S5). These results indicate that cohesin-independent interactions are likely maintained by mechanisms that do not directly depend on intact Pol II or transcription.

Taken together, these analyses demonstrate that cohesin-dependent interactions are typically longer-range and associate with the binding of cohesin and CTCF *in cis*, while cohesin-independent interactions are shorter-range and associate with features of active promoters and enhancers.

### Cohesin-dependent promoter interactions participate in transcriptional control

To address the effects of rapid depletion of cohesin on gene expression, we generated profiles of newly synthesized mRNA using SLAM-seq (Herzog et al., 2017; Muhar et al., 2018) in SCC1-AID cells treated with auxin for 60 min and untreated controls (see Methods for details). The majority of genes were relatively unperturbed upon cohesin depletion, consistent with previous findings in other cell types in “steady-state” conditions (Haarhuis et al., 2017; Nora et al., 2017; Rao et al., 2017; Schwarzer et al., 2017; Seitan et al., 2013; Sofueva et al., 2013; Wutz et al., 2017). However, we detected 421 and 266 genes whose levels of newly synthesised mRNA were significantly upregulated and downregulated in cohesin-depleted cells, respectively (adjusted p-value < 0.05; absolute log_2_ fold-change > 0.1). We also defined a set of 1,197 “constant” genes showing strong expression levels (top 25% RNA-seq signal) and no significant changes in nascent transcription upon cohesin depletion (see Methods for details; Fig. 5A). GO term enrichment analysis revealed that downregulated genes show involvement in translational processes, mRNA to ER targeting and negative regulation of gene expression. The up-regulated genes show involvement in cell-cycle processes and negative regulation of gene expression (Table S1).

**Figure 5.**
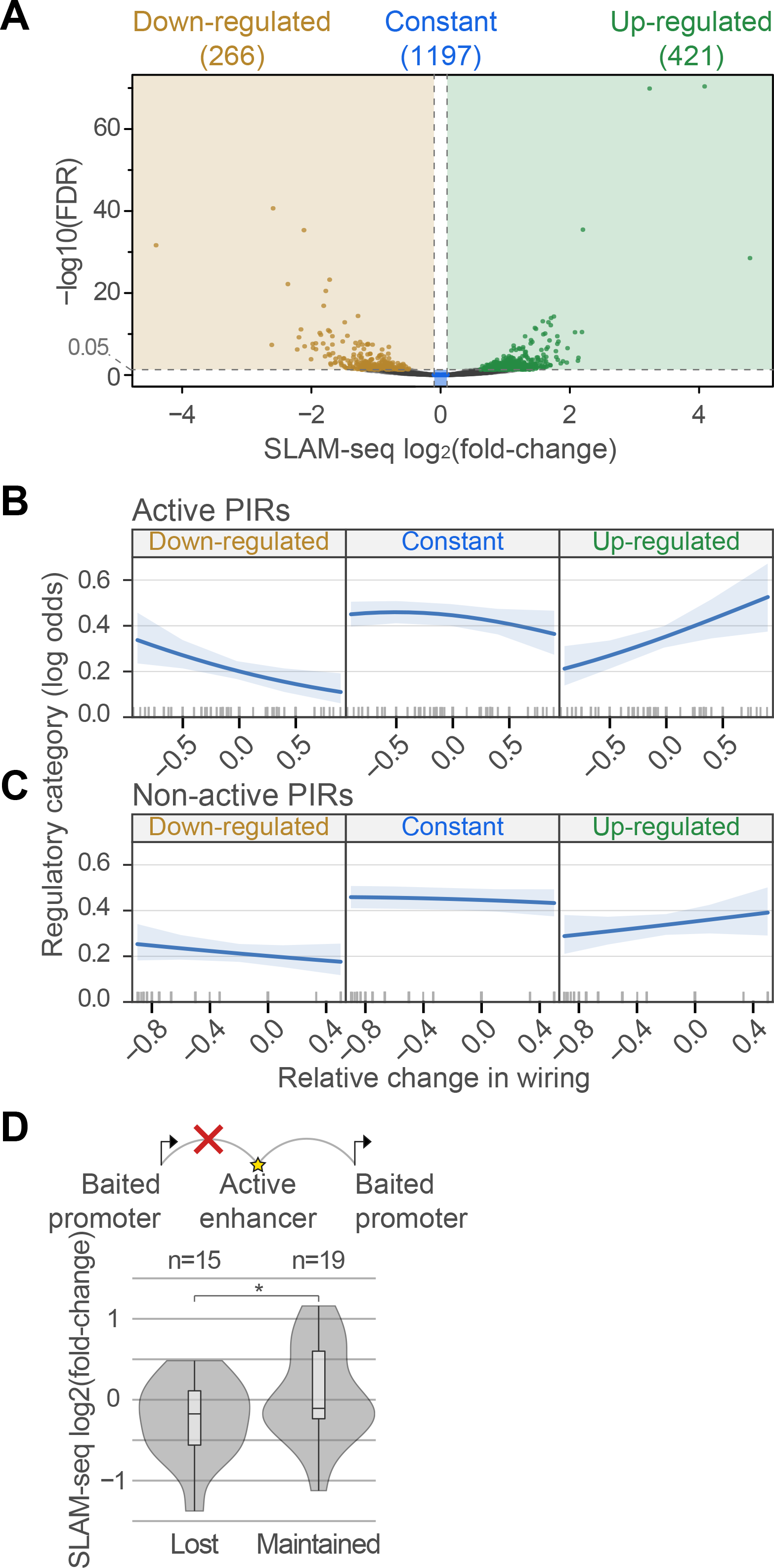
Transcriptional response upon cohesin depletion associates with rewiring of promoter interactions with active PIRs. (A) Volcano plot of SLAM-seq nascent RNA levels upon SCC1 depletion. In total, transcripts were detected for 7,424 genes, of which 266 were categorised as down-regulated and 421 as up-regulated; additionally, a subset of 1,197 highly-transcribed constant genes was identified and used in further analyses (see Methods for details). (B) Effect plot of ordinal logistic regression analysis probing an association of nascent transcriptional response to SCC1 depletion with rewiring of promoter interactions with active enhancers for the respective genes. The response variable describes gene category with respect to dynamics of nascent transcriptional dynamics upon SCC1 (see panel A). The predictor variable describes the difference between the number of gained and lost promoter interactions per bait as a proportion of all interactions per bait (see Methods for details). (C) Same as (B) but using the rewiring of PIRs not containing active enhancer annotations as the response variable. (D) Violin plots showing a log_2_ fold-change of the SLAM-seq signal upon SCC1 depletion for a small subset of genes whose promoters contact a shared active PIR in unperturbed cells, but either lose or maintain interactions with it upon SCC1 depletion. * One-sided t-test = 0.048.

We asked whether the observed changes in transcription upon cohesin depletion were consistent with the dynamics of enhancer-promoter interactions in these conditions. We devised an ordinal logistic regression model, using the nascent transcriptional response of a gene (“upregulated”, “constant”, “downregulated”) as the outcome variable, and the relative change in the numbers of connected active enhancers as the explanatory variable (see Methods for details). The direction of change in nascent transcription significantly associated with relative changes in the numbers of connected active enhancers (p = 0.02, effect size = 0.74; Figs. 5B and S4). Notably, the same was not true when PIRs without active enhancer annotations were used instead of active PIRs in the same regression framework (p = 0.59, Fig. 5C). Next, we focused on 15 active enhancers that engaged both in lost (n=15) and maintained (n=19) promoter interactions with different genes upon cohesin depletion. Transcription of genes that lost interactions with these enhancers was downregulated in cohesin-depleted cells compared with the genes that maintained interactions with the same enhancers (one-sided t-test p = 0.048, Fig. 5D). Jointly, these results suggest that rewiring of connections between active promoters and enhancers contributes towards changes in gene expression upon cohesin depletion.

Finally, to validate the causal effect of enhancers with cohesin-dependent promoter connections on target gene expression, we selected two genes, *NUAK1* and *SLC16A6*, that show abundant loss of enhancer-promoter interactions upon cohesin depletion (Fig. 6A and 1A). Previous studies have established that recruitment of the KRAB-repressor domain using dCas9 can inhibit short- and long-range enhancer function (Gilbert et al., 2013; Thakore et al., 2015). Inhibiting the enhancers of these genes located at cohesin-dependent PIRs by targeting dCas9-KRAB to them (in a pooled setting) resulted in a significant downregulation of both of these genes (Fig. 6B). These observations support the model that a subset of cohesin-dependent interactions connect promoters to functionally relevant enhancers.

**Figure 6.**
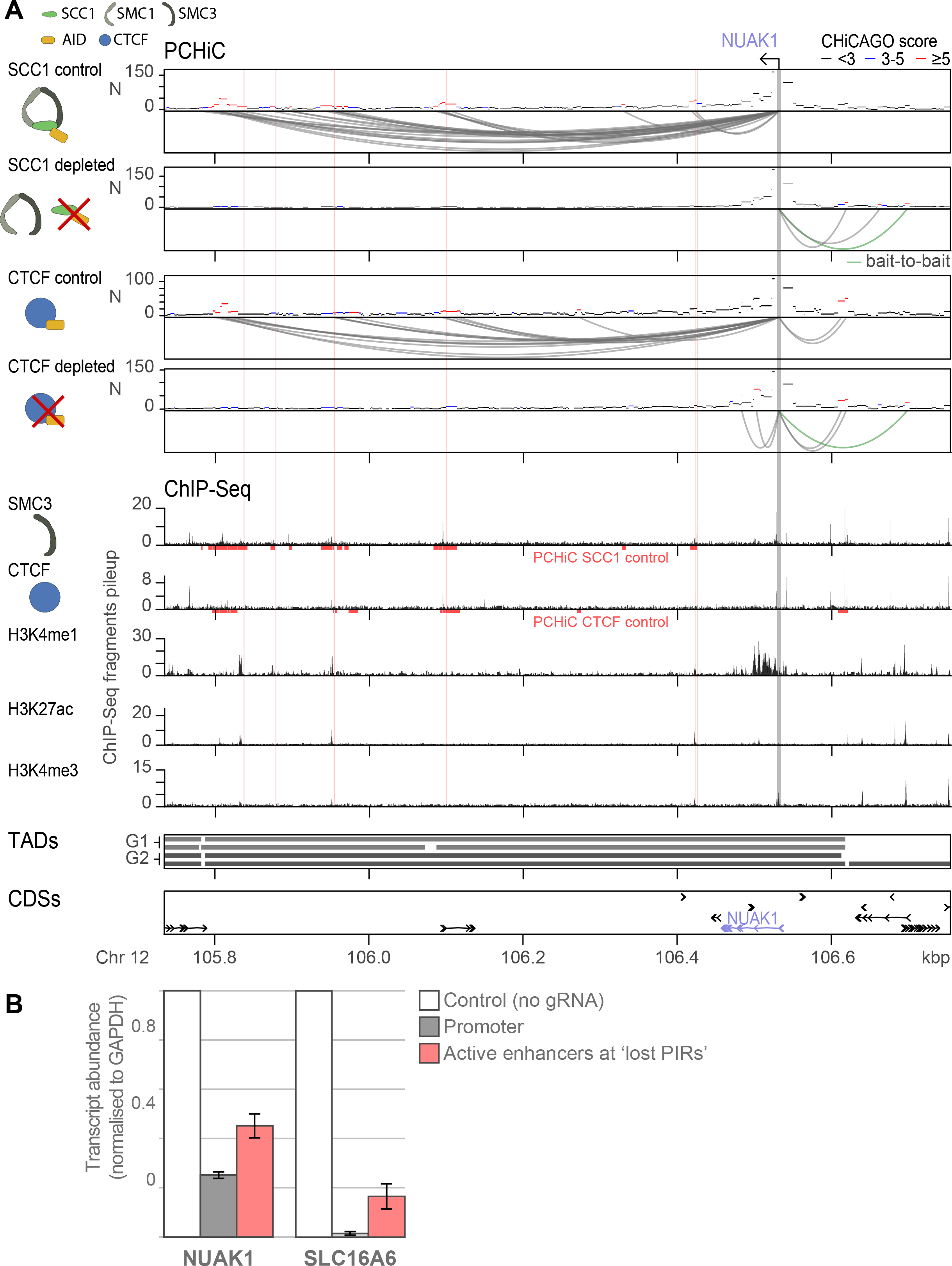
Perturbation of enhancers within cohesin-dependent PIRs affect target gene expression. (A) Chromosomal interactions detected for *NUAK1* promoter by PCHi-C. The top four tracks show PCHiC interaction profiles with interaction arcs for the conditions: SCC1 control, SCC1 depleted, CTCF control, CTCF depleted. The following five tracks show ChIP-seq pileups in interphase for the targets: SMC3, CTCF, H3K4me1, H3K27ac, H3K4me3. The bottom two tracks show TAD intervals in cell cycle stages G1 and G2, and coding sequences (GRCh37). Upon depletion of SCC1 or CTCF, *NUAK1* promoter loses the majority of its promoter interactions. Enhancers that contact *NUAK1* in a cohesin-dependent manner were selected for dCAS9-KRAB based on the presence of H3K4me1 and H3K27ac. (B) Bar chart showing relative transcript abundance (by RT-qPCR) upon targeting dCAS9-KRAB to the promoter regions of *SLC16A6* or *NUAK1* (grey bars) and separately to the selected active enhancers (in a pooled setting).

## Discussion

In this study we have used high-resolution Promoter Capture Hi-C in conjunction with nascent transcript sequencing by SLAM-seq to study the effects of rapid depletion of cohesin and CTCF on promoter interactions. We report extensive rewiring of promoter interactions upon perturbation of these architectural proteins, which associate with changes in target gene transcription. Loss of large numbers of promoter-enhancer interactions upon cohesin depletion is consistent with superenhancer rewiring observed with Hi-C in HCT-116 cells (Rao et al., 2017), as well as with recent observations from a Promoter Capture Hi-C experiment in this cell line (El Khattabi et al., 2019). We find that the strongest predictors of cohesin dependence of PIRs are interaction distance and the binding of architectural proteins *in cis*. This observation confirms the long-standing reports of architectural protein binding *in cis* to promoters and enhancers (Dorsett and Merkenschlager, 2013; Yan et al., 2018), and is line with recent findings that deletion of CTCF motifs can disrupt enhancer-promoter interactions within TADs (Ren et al., 2017). These observations reinforce the importance of the effects of architectural protein binding *in cis* to regulatory elements in gene expression control. However, we also detect a smaller subset of generally short-range promoter interactions that emerge *de novo* upon architectural protein depletion, which often cross native TAD boundaries. Collectively, these results indicate that cohesin and CTCF have two important and separable functions in gene regulation: promote long-range interactions between promoters and regulatory elements and insulate promoters from enhancers in neighbouring TADs.

Importantly, we find that a significant minority of promoter interactions are retained upon architectural protein depletion. We find that these interactions often connect active promoters with active promoters and enhancers. Notably, most of these interactions are retained throughout interphase, and a subset are also observed in mitosis, in contrast to the largely transient interactions between *cis-*regulatory regions observed in early G1 that may reflect the A/B compartment signal (Zhang et al., 2019). This finding is consistent with observations that cohesin depletion abrogates TADs but not all chromosomal interactions (El Khattabi et al., 2019; Gassler et al., 2017; Rao et al., 2017; Schwarzer et al., 2017; Wutz et al., 2017)is also in line with evidence of cohesin-independent fine-scale chromosomal topology obtained in recent microscopy studies (Bintu et al., 2018; Luppino et al., 2019). These cohesin-independent contacts are presumably separate in nature from the well-characterised A/B compartment structure that is also independent of architectural proteins (Nora et al., 2017; Nuebler et al., 2018; Rao et al., 2017; Wutz et al., 2017).

The PIRs of cohesin/CTCF-independent interactions are enriched for active histone modifications and recruit chromatin-binding co-factors such as p300 acetyltransferase and BET family protein BRD4. The binding of p300 is a canonical hallmark of active enhancers (Heintzman et al., 2007; Visel et al., 2009), and BRD4 is also known to bind active chromatin regions (Kanno et al., 2014; Lovén et al., 2013; Zhang et al., 2012). BRD4 was previously shown to play a role in chromatin insulation, while depletion of BRD4 led to widespread chromatin decompaction (Floyd et al., 2013; Wang et al., 2012). A role in chromatin insulation was also previously reported for another BET family protein, BRD2, notably in a CTCF-dependent manner (Hsu et al., 2017). Other factors associated with cohesin-independent interactions include chromatin regulatory proteins such as NuRD-related factor PWWP2A, DREAM complex subunit LIN-9 and histone demethylase PHF8, which are known to be involved in the transcriptional control of the cell cycle (Liu et al., 2010; Pünzeler et al., 2017; Sadasivam and DeCaprio, 2013). Both PWWP2A and LIN-9 associate with H2A.Z histone variant (Latorre et al., 2015; Pünzeler et al., 2017), while interestingly, the DREAM complex was found to possess insulator activity in *Drosophila* (Bohla et al., 2014; Korenjak et al., 2014). Finally, an enrichment for H3K27me3 histone mark associated with Polycomb Repressive Complexes is consistent with the independent findings of a very recent study in ES cells (Rhodes et al., 2020). Notably, the enrichment of Polycomb at the PIRs of cohesin-independent interactions appears to be considerably more pronounced in ES cells compared with differentiated cells such as HeLa (Rhodes et al., 2020), in line with the prominent role of these complexes in stem cell genome architecture and transcriptional control (Mas and Di Croce, 2016; Schoenfelder et al., 2015b).

The broad range of chromatin co-factors detected at the PIRs of cohesin-independent interactions points to the potential diversity of mechanisms that could support promoter interactions independently of cohesin and CTCF. Our results suggest that these interactions are also generally independent of RNA Pol II and transcription, and a recent study using a different RNA Pol II inhibitor has independently arrived at the same conclusions (El Khattabi et al., 2019). The enrichment for promoter-promoter contacts points to the possibility that some cohesin-independent interactions reflect phase-separated “transcription factories”, which can be stabilised by intrinsically disordered proteins including the Mediator complex and BRD4. The Mediator complex has long been proposed to physically link enhancers and promoters (Malik and Roeder, 2010), but deletion of its core subunit MED14 did not induce major changes in promoter-enhancer interactions (El Khattabi et al., 2019). Other potential candidates include the LDB1 complex that has been shown to facilitate promoter-enhancer interactions in some systems (Deng et al., 2012; Krivega and Dean, 2017; Song et al., 2007), as well as eRNAs that may be involved in chromatin loop stabilisation (Matharu and Ahituv, 2015).

Although the binding of cohesin and CTCF is not restricted to enhancers, we confirm that these proteins mediate subsets of promoter-enhancer interactions that are relevant for gene expression, consistent with previous reports (Merkenschlager and Nora, 2016; Seitan et al., 2013; Xu et al., 2016). However, the fact that many interactions with active enhancers are retained upon architectural protein depletion, and only a small number of contacts are gained in these conditions, may explain why this perturbation does not have a more pronounced effect on steady-state transcription. How can these observations be reconciled with the requirement for cohesin for gene induction (Cuartero et al., 2018) and lineage-specific gene regulation (Liu et al., 2019)? One possibility is that in non-transcriptionally-permissive chromatin contexts, cohesin/CTCF-independent promoter contacts are unable to form in the absence of some prior activation events, such as chromatin decompaction of the locus or establishment of appropriate chromosomal conformation. It is possible that these events require cohesin-dependent promoter interactions to act as “pioneers” (Darbellay and Duboule, 2016). This model is broadly consistent with the findings that the first enhancer in chains of enhancers is often distal and contains CTCF motifs (Song et al., 2019). It may also explain phenomena, whereby a weak distal enhancer, but not a strong proximal one, is required for developmental activation of sex-determining gene *Sox9 (Gonen et al., 2018)*. Elucidating the interplay of cohesin-dependent and independent mechanisms of long-range gene control is an important research direction for future studies, with broad implications for health and disease.

## Materials and Methods

### Cell culture

HeLa Kyoto cells were used as a model system for all experimental analyses. Cells were cultured at 37 in suspension in DMEM supplemented with 10% FCS, glutamate and penicillin-streptavidin.

### Cell cycle synchronisation

HeLa cells were synchronised at the G1/S-phase transition by two consecutive cell cycle arrest phases using 2mM thymidine and released into fresh medium for 6h (G2-phase), or 15h (G1-phase). For mitotic cells, Nocodazole (100 ng/ml) was added 8h after release from double thymidine block, to arrest the cells in prometaphase. Post-mitotic cells were removed by shake off after five hours.

### WAPL/PDS5A/PDS5B RNA interference

HeLa cells were transfected with siRNAs as described in (Lelij et al., 2014) followed by addition of thymidine. The siRNA sequences directed against WAPL, PDS5A and PDS5B (Table S2) were obtained from Ambion. Transfection was performed by incubating duplex siRNA with RNAi-MAX reagent (100 nM) in growth medium lacking antibiotics. Cells were harvested after 72h of RNAi treatment.

### Auxin induced degradation of SCC1 and CTCF

Auxin-inducible degron systems for the analysis were generated as described in (Wutz et al., 2017). Briefly, C-terminal tagging of *SCC1* and *CTCF* with an Auxin induced degron (AID) was performed using CRISPR/Cas9-mediated genome editing with a double-nicking approach (Ran et al., 2013). The C-termini were extended with monomeric EGFP (L221K) and the IAA1771-114 (AID*) mini-degron from *Arabidopsis thaliana* (Morawska and Ulrich, 2013). Single clones were selected using flow cytometry PCR was used to confirm that all alleles were successfully modified. Cells stably expressing Tir1 for auxin-inducible protein degradation were generated by transduction of SCC1-AID and CTCF-AID knock-in cells with the lentiviral vector SFFV-OsTir1-3xMyc-T2A-Puro (SOP)(Wutz et al., 2017) and Puromycin-selection (2 µg/ml, Invitrogen).

### Promoter Capture Hi-C (PCHi-C)

PCHi-C was performed on SCC1-mEGFP-AID and CTCF-mEGFP-AID HeLa cells synchronized in G1 and treated with auxin for either 0 or 120 minutes, WAPL/PDS5A/PDS5B RNAi knockdown HeLa cells, as well as unmodified HeLa cells synchronized in G1, G2 or mitosis. *HindIII*-based Hi-C libraries from our previous study (Wutz et al., 2017) were captured using SureSelect target enrichment system (Agilent Technologies) according to manufacturer’s instructions, using custom-designed RNA bait library and paired-end blockers. Promoter-based ligated products were isolated and amplified as previously described (Schoenfelder et al., 2018) and paired-end sequenced using Illumina HiSeq 2500 at the Babraham Institute Sequencing Facility.

### SLAM-seq

Transcriptional responses to SCC1-degradation were profiled by SLAM-seq as described previously (Muhar et al., 2018) with minor modifications. HeLa SCC1-AID cells were synchronized in G1 by double thymidine block as for PC-HiC. SCC1 degradation was induced by addition of 500 µM auxin (indole-3-acetic acid sodium salt, Sigma-Aldrich). Newly synthesized RNA was labelled by addition of 4-Thiouridine (4SU, Carbosynth) at a final concentration of 100 µM for 60 minutes. Cells were snap-frozen and extracted total RNA was subjected to alkylation by iodoacetamide (Sigma-Aldrich) for 15 minutes at room temperature in an appropriate buffer. Alkylation was stopped by addition of dithiothreitol (DTT, GE-Healthcare) and alkylated RNA purified by ethanol precipitation. 3’ end mRNA sequencing libraries were generated from 500 ng of alkylated RNA using the QuantSeq 3’ mRNA-Seq Library Prep Kit FWD for Illumina (Lexogen). Single-read sequencing was performed by the Vienna Biocenter Core Facilities (VBCF) for 100 cycles on a HiSeq2500 sequencer (Illumina).

### Triptolide treatment and Western blot

Scc1-EGFP-AID cells were treated with 300 nM Triptolide (Sigma, T3652) for 3 hours, and cells were collected for western blot or Hi-C. For western blot, cells were resuspended in RIPA buffer (50 mM Tris pH 7.5, 150 mM NaCl, 1 mM EDTA, 1% NP-40, 0.5% Na-deoxycholate and 0.1% SDS), which additionally contained pepstatin, leupeptin and chymostatin (10 µg/ml each) and PMSF (1 mM). The protein concentration was determined with the Bradford Protein Assay (Bio-Rad Laboratories). SDS-PAGE and standard Western blot technique were applied to detect individual proteins with specific antibodies. Rabbit anti-RNA polymerase II Ser2 (abcam, ab5095), rabbit anti-RNA polymerase II Ser5 (abcam, ab5131) and mouse anti-tubulin (Sigma, T-5168).

### CRISPRi using dCas9-KRAB

HeLa SCC1-AID cells stably expressing dCas9-KRAB for CRISPRi-mediated repression of enhancer or promoter activity were generated by transduction with the lentiviral vector pHR-SFFV-KRAB-dCas9-P2A-mCherry (Gilbert et al., 2014) (Addgene plasmid #60954) and subsequent sorting for mCherry-positive cells using a FACS Aria III cell sorter (BD Lifesciences). Lentiviral packaging was performed in Lenti-X packaging cells (Clontech) transfected with lentiviral transfer plasmids and packaging plasmids helper plasmids pCMV-VSV-G (Addgene plasmid #8454) and pCMVR8.74 (Addgene plasmid #22036) using standard procedures. gRNA oligos (sequences in Table S3A) were cloned into the expression vector (JJ4_sgRNA-U6-IT-EF1As-Thy1.1-P2A-NeoR) and transfected into packaging cells (293FT, Thermo Fisher, R70007) using Fugene 6 (Promega, E2691). Virus was then harvested and applied to Scc1-EGFP-AID/dCas9-KRAB cells. Infected cells were selected by G418 (Gibco, 108321-42-2) and used for RT-qPCR.

### RT-qPCR

RNA was prepared using TRIzol (Thermo Fisher, 15596026) according to the manufacturer’s instructions. The cDNA was generated by reverse transcription using Random hexamers (Thermo Fisher, N8080127) and SuperScript™ II Reverse Transcriptase (Thermo Fisher, 18064014) according to manufacturer’s instructions. Different transcripts were then compared by qPCR using GoTaq® qPCR Master Mix (Promega, A6001) and primers specific for selected genes and GAPDH (see Table S3B). *GAPDH* levels were used for normalization of the selected genes and data were presented as fold change compared to the control gRNA.

### Definition of topologically associating domains

Quality-controlled, aligned and filtered Hi-C datasets (Wutz et al., 2017) were processed with HOMER (Heinz et al., 2010). Hi-C reads were binned at a 40kb resolution and normalised using iterative correction (Imakaev et al., 2012). Directionality index (DI) scores (Dixon et al., 2012) were calculated using a 5kb step size, a 25kb window size and a 1Mb upstream and downstream window size, and were subsequently smoothed using 25kb kernel density smoothing. Topologically associating domain (TAD) boundaries were called as local extrema in DI transitions from negative to positive were detected. TADs were calculated by setting *minIndex = 1* and *minDelta = 2*.

### PCHi-C processing and analysis

Promoter interactions were called using the CHiCAGO pipeline (version 1.1.5) (Cairns et al., 2016). CHiCAGO models expected promoter interaction frequencies based on the Delaporte distribution, which has a negative binomial component for random Brownian collisions and a Poisson component for technical noise. CHiCAGO corrects for multiple testing by means of a p-value weighting procedure based on the expected true positive rate at a given interaction distance estimated on the basis of consistency between biological replicates. CHiCAGO scores represent soft-thresholded -log weighted p-values. Enrichment of multiple chromatin marks is typically maximised at promoter interacting regions (PIRs) at a CHiCAGO score of 5 (Cairns et al. 2016; Freire Pritchett et al. eLife 2017) and this score cutoff was used to call significant promoter interactions.

### Clustering of promoter interactions

K-means clustering was used to partition promoter interaction scores across samples. The function *kmeans* from the base R package *stats* was called with the Hartigan-Wong clustering algorithm, a maximum of 1000 iterations and 25 initial random configurations. Prior to clustering, CHiCAGO scores were asinh-transformed and capped at median + 3 MAD. The number of clusters k was determined by iteratively performing k-means clustering with k-values ranging from 2 to 16 and choosing a partitioning with a low k as well as a low total within-cluster sum of squares value. Additionally, partitionings that resulted in multiple clusters with similar centroid values were avoided. Using these criteria, a partitioning with k = 13 was selected.

### Differential PCHi-C interaction calling

Chicdiff (Cairns et al., 2019) was used to detect differential promoter interactions between SCC1-AID control (Aux-) and depleted (Aux+) samples, and between CTCF-AID control (Aux-) and depleted (Aux+) samples. Chicdiff uses DESeq2 (Love et al., 2014) differential interaction calling with custom feature selection, normalisation and multiple testing correction procedures. Testing was performed on promoter interactions exceeding a Chicago score cutoff of 5 detected on merged-replicate data for either of the two conditions. Chicdiff v0.2 was used, using the normalisation procedure available in the later versions as *norm = “fullmean”*. The difference in the mean asinh-transformed Chicago scores between conditions above 1 was used to prioritise the potential “driver” differential PIRs within the tested aggregated fragment bins, which is the default setting.

### Defining promoter interaction rewiring categories upon SCC1 depletion

We constructed consensus sets of interactions that are lost, maintained and gained upon SCC1 depletion based on results from Chicdiff analysis, Chicago promoter interaction calling for each condition and K-means clustering. The criteria were as follows. “Lost”: Chicdiff adjusted weighted p-value ≤ 0.01; log2 fold-change < 0; K-means clusters B, C, D or F; “gained”: Chicdiff adjusted weighted p-value ≤ 0.01; log2 fold-change > 0; K-means cluster K; “maintained”: Chicdiff adjusted weighted p-value > 0.01; Chicago score ≥ 5 in the merged-replicate samples for each condition; difference in the mean asinh-transformed Chicago scores between conditions not exceeding 1; K-means clusters A, E or J). The absolute majority (>87%) of Chicdiff-detected differential interactions had consistent K-means cluster assignment (36,174/40,666 lost and 2,484/3,653 gained, respectively).

### Analysis of the distribution of promoter interactions within TADs

To create the diagrams in Figure 3A, the linear genomic distance between a given restriction fragment (RF; either the baited promoter fragment or PIR) and the nearest TAD boundary was normalised to the length of the TAD:

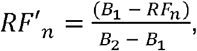

where RF’_n_ is the relative location of the n^th^ restriction fragment with respect to the nearest TAD boundary; B_1_ and B_2_ are the start and end of the nearest TAD, respectively; and RF_n_ is the centre coordinate of the n^th^ restriction fragment. RF’_n_ takes the value of 0 when the fragment is located at the TAD start boundary and the value of 1 when the RF is located at the TAD end boundary.

RF’_n_ was calculated for each RF, and the frequency distribution of RF_1-N_ was then calculated and visualised for the interval: RF’_n_ = [-2, 3]. Note that this interval corresponds to the x-axes of the panels in Figure 3A, showing the values [-2, -1, S, E, 1, 2]. The frequency distributions for multiple TAD partitionings were combined by calculating the mean and the standard error per bin.

RF’_n_ distributions were plotted separately depending on the relative location of the second interaction partner (i.e., either the baited promoter or PIR) within a pentile interval with respect to TAD length (TAD boundary-proximal: 0-20% and 80-100%; intermediate: 20-40% and 60-80%; mid-TAD: 40-60%). The results for the boundary-proximal and intermediate windows were combined by taking the reflection along the TAD centre of the frequency distributions.

### TAD window enrichment analysis

RF enrichment within TAD windows (Figure 3B) was performed by dividing the observed proportion of RFs in a pentile window (peripheral, intermediate and central) by the expected proportion according to the uniform distribution. For the central window the expected proportion is 0.2 (one fifth), while for peripheral windows, as well as for intermediate windows, the expected proportion is 0.4 (since there are two of each of these windows). Enrichment is then defined as:

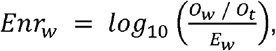

where *Enr_w_* is the enrichment in window w (peripheral, intermediate or central), *O_w_* is the observed number of interacting RFs in window w, *O_t_* is the observed total number of interacting RFs in the TAD, and *E_w_* is the expected proportion of RFs under the uniform distribution for window w (0.4, 0.4, or 0.2 respectively). The enrichment scores are calculated for TAD partitionings obtained on each of the four datasets (two replicates of G1- and G2-synchronised HeLa cells, respectively) and the mean +/-1SE are shown in Figure 3B.

### Association between cohesin dependence and TAD boundary crossing of promoter interactions

For the analysis in Figure S2, logistic regression was performed at the level of individual promoter interactions, where the Boolean dependent variable was set to 1 if a promoter interaction crosses a TAD boundary in all four TAD partitionings (generated from two replicates of G1- and G2-synchronised HeLa cells, respectively), and to 0 otherwise. Promoter interactions crossing TAD boundaries defined in a subset of partitioning were removed from the analysis. The independent variables were the log_10_(bp) promoter interaction distance and the rewiring category (lost, maintained or gained). A ‘sum to zero’ contrast matrix was used. The R function *allEffects* from the *effects* package (Fox, 2003) was used for visualisation.

### ChIP-seq data sources and processing

HeLa-specific ChIP-Seq data were obtained from two sources: the ENCODE project (ENCODE Project Consortium, 2012) and the Gene Expression Omnibus (GEO) (Edgar, 2002). ENCODE files were downloaded manually as BAM files, aligned against GRCh37. Dataset and publication IDs are listed in Table S4. Quality control on ENCODE files was performed using the “Read coverage audits” QC metrics provided on the ENCODE website (https://www.encodeproject.org/data-standards/audits/). Only entries with replicate data and with a read length of ≥ 36 were included and files with read-depth limitations or complexity limitations were excluded (i.e. the dataset must have ≥ 10 million usable fragments and at least “moderate library complexity” is required). Separately, unaligned data were downloaded from GEO as fastq files using the sra_fqdump command in Cluster Flow (Ewels et al., 2016). Read quality was tested using FastQC (http://www.bioinformatics.babraham.ac.uk/projects/fastqc/), and FastQscreen (Wingett and Andrews, 2018) was used to test for contaminants and to ascertain that the sequencing material indeed originated from the human genome. Overall, 19 ChIP-Seq targets were excluded from further analysis. Datasets passing the QC were processed with Trim Galore (https://www.bioinformatics.babraham.ac.uk/projects/trim_galore/) to remove adapters. Reads were then aligned against GRCh37 using Bowtie2 (Langmead and Salzberg, 2012)

### Definition of ChIP-seq signal enrichment per restriction fragment

Read counts per restriction fragment (RF) were obtained by using htseq-count (Anders et al., 2015), requiring reads to have a minimum mapping score of 10 and with the -m flag set to ‘union’. RFs that were shorter than 50bp or longer than 50kbp were excluded from further analysis. The Anscombe variance-stabilising transformation was applied to RF-level read counts (Harrison, 2017), with the dispersion parameter set to 0.4. Following that, between-chromosome quantile normalisation was applied (separately for each ChIP-Seq dataset) using the *normalizeQuantile* function from the *aroma.light* R package (Bengtsson et al., 2004). Linear regression was then used to define the relationship between the Anscombe-transformed, quantile-normalised RF-level read counts (response variable) and the RFs’ log_10_-length (predictor variable). Replicates were merged by taking the mean residual score per RF. A cutoff on the studentized residuals from these regression models was then applied to define RFs with enriched signals in each dataset. The cutoff was taken to be 1.6449, corresponding to Student distribution p-value = 0.05 at df = n-k-1, where n = 801363 (the number of RFs in the model) and k=1 (the number of explanatory variables in each model).

### Features of PIRs associated with maintained versus lost interactions

The analysis was performed using LASSO logistic regression, with the rewiring category (lost versus maintained in the absence of cohesin) as the response variable. The predictor variables were the binary ChIP-seq signal enrichment scores for each analysed dataset computed as above, asinh-transformed Chicago interaction scores, log10-transformed linear distance between interaction partners, and a flag indicating promoter-promoter interactions. The distance and interaction score parameters were scaled and centred to mean = 0 and SD = 1. Since a PIR can have multiple interactions, we performed regressions separately using the lowest, mean (Figure 4D) and highest score per PIR and the distance of the respective interaction, which produced comparable results (data not shown). PIRs associated with interactions from both categories (i.e. lost as well as maintained) were excluded from this analysis.

The *glmnet* function from the glmnet R package (Friedman et al., 2010) was used to perform LASSO logistic regression, with the *alpha* flag set to 1 (indicating LASSO regression) and the *thresh* parameter set to 10^-12^. Cross-validation was performed using the function *cv.glmnet*. Significance was tested at a λ value that minimises the number of parameters while remaining within 1SE of the cross-validated error (λ 1SE). To infer p-values on the regression coefficients, the function *fixedLassoInf* from the R package *selectiveInference* (Lee et al., 2016) was used with parameters *tol.beta* = 10-3-, and *tol.kkt* = 0.3.

### Association between the dynamics of promoter-enhancer interactions and nascent transcription upon cohesin depletion

#### Ordinal logistic regression approach

Dynamics of nascent transcription based on SLAM-seq data was used as the outcome variable and the relative change in the number of promoter interactions upon cohesin depletion as the predictor variable. SLAM-seq data were processed using the SLAM-DUNK pipeline (Neumann et al., 2019).

The relative change in the number of interactions per baited promoter was computed as follows:

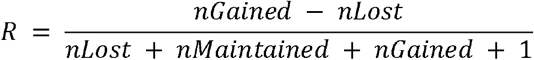

where *nGained*, *nMaintained* and *nlost* epresent the numbers of gained, lost and maintained promoter interactions per baited fragment, respectively, and a pseudocount of 1 is added to avoid division by 0. This quantity was computed separately for promoter interactions with PIRs (*R*_active_) containing active enhancers and for promoter interactions with other PIRs (*R*_non-active_) and used in the analyses shown in Figures 5B and 5C, respectively. To define PIRs containing active enhancers, we used the studentized residuals from the regression analysis to determine ChIP-seq signal enrichment. PIRs with top 5% studentized residuals for H3K4me1 and H3K27ac or H3K4me3 were considered as containing active enhancers.

To construct the outcome variable, genes were assigned into three categories (downregulated, constant, upregulated; Figure 5A), based on the log2 fold-change (LFC) in SLAM-seq read count and DESeq2 FDR-adjusted p-value between the control and SCC1-depleted samples (downregulated: LFC ≤-0.1 and p-value ≤0.05; constant: |LFC| ≥ 0.1, p-value > 0.05, RNA-seq RPM > 0.21 (75th percentile); upregulated: LFC ≥0.1, p-value ≤ 0.05). Genes not meeting the criteria for either of the three categories were removed from the analysis. Ordinal logistic regression was then performed using the polr function from the MASS package (Venables and Ripley, 2002). The proportional odds assumption was tested by performing a graphical parallel slopes test (data not shown).

#### Permutation-based approach

Contingency tables for association between transcriptional regulation and promoter interaction rewiring upon SCC1 depletion were constructed for lost, maintained, and gained promoter interactions separately (see Table S### for an illustrative contingency matrix). Promoter interaction rewiring and transcriptional response were then expressed as a log odds ratio:

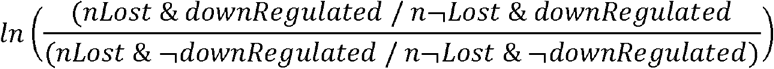

This ratio is > 0 if genes with lost promoter interactions tend to be more strongly down-regulated than genes with maintained or gained promoter interactions. Note that these log odds ratios were calculated for all combinations of rewiring category and regulatory category (e.g., lost & up-regulated, lost & non-regulated, maintained & down-regulated, etc). To construct regulatory categories (downregulated, constant and upregulated), a more stringent log2 fold change cutoff was applied (downregulated: LFC ≤0.1 and p-value ≤0.05; constant: |LFC| ≥0.1 and p-value > 0.05, upregulated: LFC ≥ 0.1, p-value ≤ 0.05). Note that in this analysis, there was no requirement for PIRs to bear active marks.

An empirical null distribution of log-odds ratios was then computed for each of these combinations, by constructing the contingency tables on randomly sampled baits (sampled from the set of baits that were categorised as described above) over 10,000 iterations. Subsequently, empirical p-values were calculated by dividing the number of times the values in the expected distribution exceeded the observed value by the total number of values in the expected distribution. The p-values were converted to z-scores (Figure S4) using the following calculation:

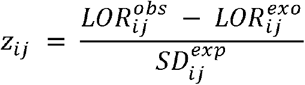

Where *z_ij_* is the z-score for the *i^th^* rewiring category (lost, maintained or gained) and the *j^th^* regulatory category (downregulated, constant or upregulated), *LOR^obs^* is the observed log_2_ odds ratio, *LOR^exp^* is the mean expected log_2_ odds ratio and *SD^exp^* is the standard deviation of the distribution of expected log odds ratios.

### Gene Ontology enrichment analysis

The online tool LAGO (https://go.princeton.edu/cgi-bin/LAGO) (Boyle et al., 2004) was used to determine GO term enrichment for genes that showed a transcriptional response upon SLAM-seq analysis. Two analyses were performed by providing LAGO with a list of gene IDs of the up-regulated genes upon SCC1 depletion and separately by providing a list of gene IDs of the down-regulated genes upon SCC1 depletion. The results were merged and similar GO terms were grouped manually. The resulting table is available in supplementary file ###.

## Data and code availability

All raw sequencing data for this project will be made available online through Gene Expression Omnibus (https://w.ncbi.nl.nih.gov/geo/). Processed datasets have been uploaded to Open Science Framework (https://osf.io/brzuc/). Scripts used to execute the analyses will be made available upon request.

## Author contributions

Conceptualization: M.J.T., G.W., M.S; formal analysis: M.J.T; funding acquisition: M.S., P.F., S.S., J.Z, J.M.P; investigation: G.W., M.M., W.T., S.B., M.J.T., V.M; methodology: G.W., M.M., W.T., V.M., M.S., S.S., P.F; project administration: M.S., G.W; resources: G.W., V.M., S.S., P.F., J.M.P., M.S.; software: M.J.T., M.S., M.M., V.M.; supervision: M.S., J.M.P., S.S., P.F., validation: W.T., M.M, G.W.; visualization: M.J.T., M.S.; writing - original draft: M.S with M.J.T.; writing – review & editing: all authors.

## Acknowledgements

The authors would like to thank Csilla Varnai, Will Orchard; Steven Wingett, Simon Andrews and Anne Segonds-Pichon (Babraham Bioinformatics Facility); Christina Tabada (Babraham Sequencing Facility) for technical assistance and advice. We thank Jonathan Cairns, Peter Rugg-Gunn, Chris Wallace, Jörg Morf, Matthias Merkenschlager and all members of our laboratories for helpful discussions. MJT was supported by a studentship from the UK’s Medical Research Council (MRC). Research in JMP group is supported by Boehringer Ingelheim, the Austrian Science Fund (FWF special research program SFB F34 “Chromosome Dynamics” and Wittgenstein award Z196-B20), the Austrian Research Promotion Agency (Headquarter grants FFG-834223 and FFG-852936) and the European Research Council (ERC) under the European Union’s Horizon 2020 research and innovation programme (Grant Agreement 693949). MS acknowledges core support from the UK’s Biotechnology and Biological Sciences Research Council (BBSRC) and MRC.

## Supplementary Figure Legends

**Figure S1.**
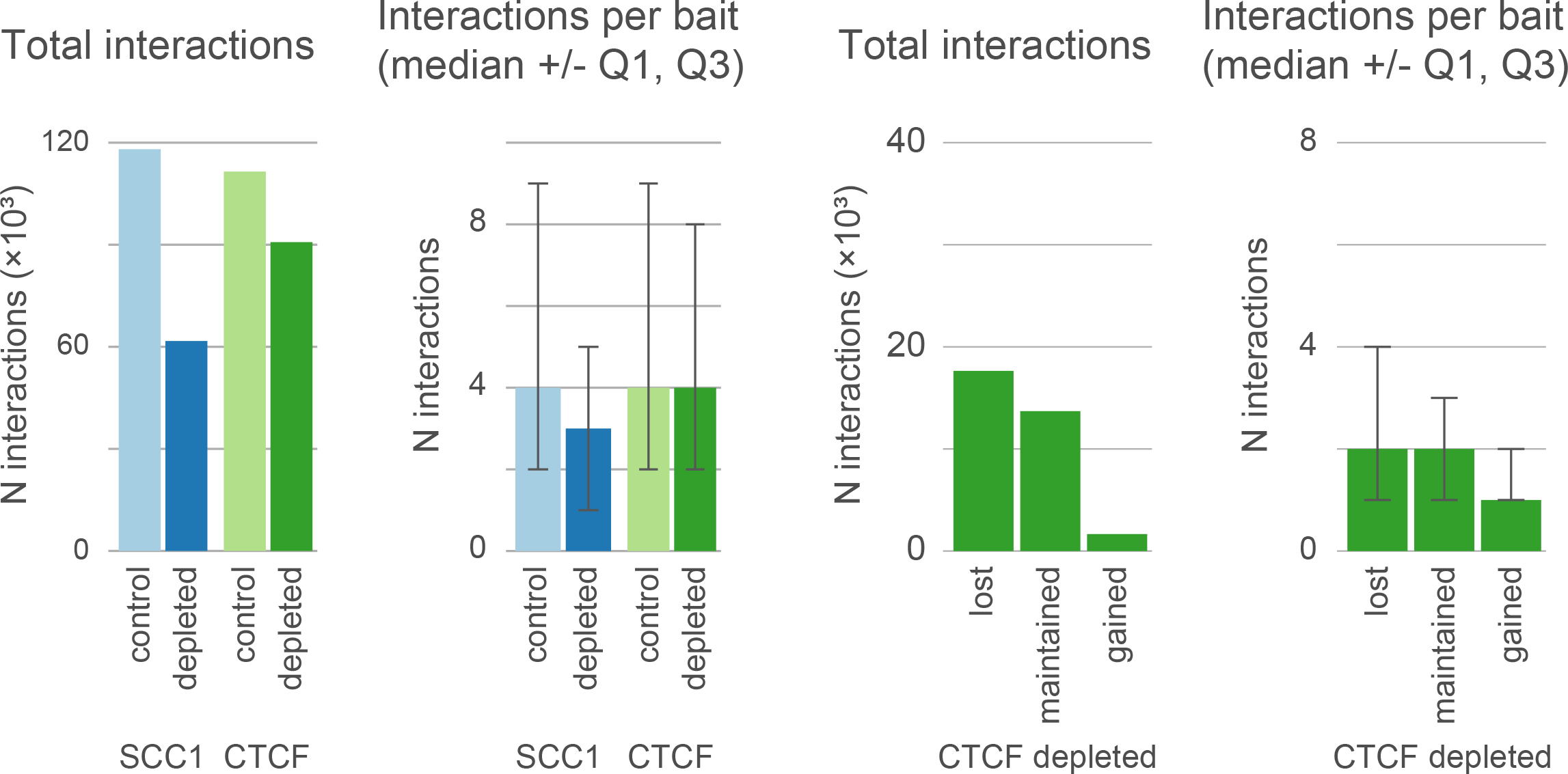
Summary statistics of promoter interactions in SCC1- and CTCF-depleted and control conditions. (A) The number of significant promoter interactions called using CHiCAGO. (B) The number of significant promoter interactions per bait. (C) The numbers of lost, maintained and gained promoter interactions upon CTCF depletion. See Figure 2A for the same statistics for SCC1 depletion.

**Figure S2.**
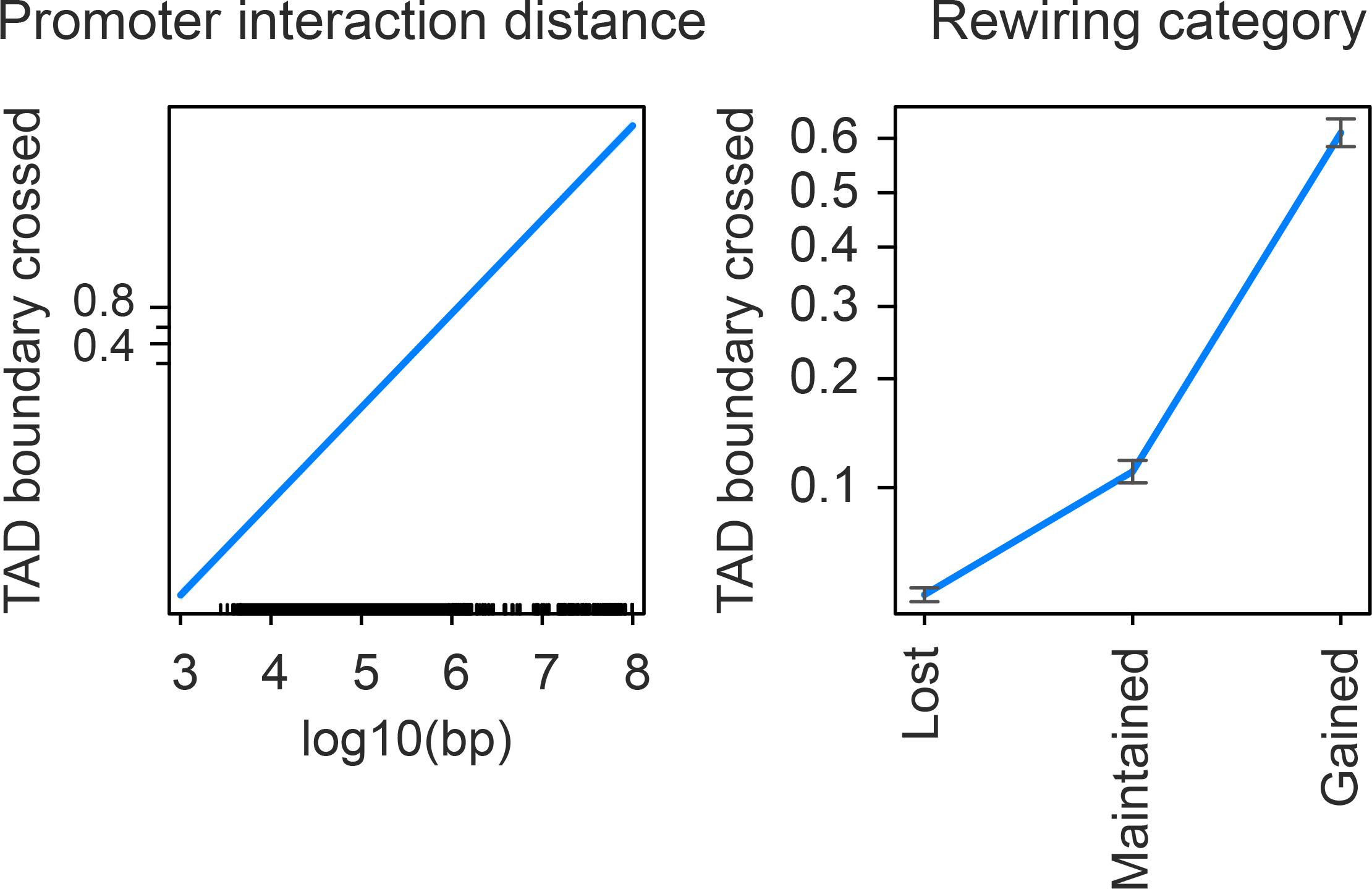
The linear distance and response to cohesin depletion associate with TAD boundary crossing of promoter interactions. Logistic regression effect plot showing the odds of TAD boundary crossing as a response variable versus log_10_ linear interaction distance (left) and interaction rewiring in response to SCC1 depletion (right).

**Figure S3.**
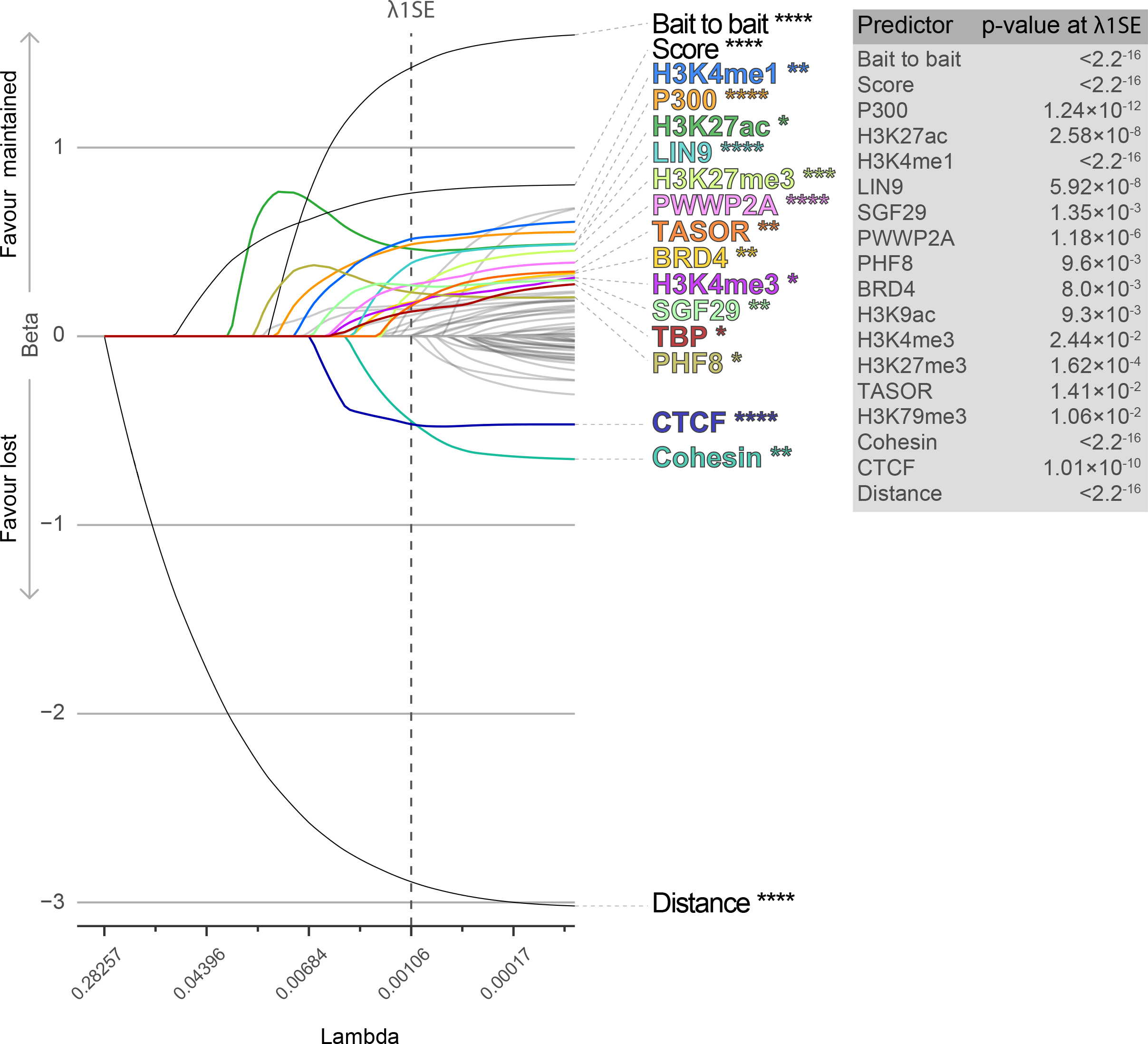
LASSO logistic regression regularisation paths for features associated with maintained versus lost interactions upon SCC1 depletion. Plots showing LASSE logistic regression coefficients (y-axis) at different values of the shrinkage parameter λ (x-axis), ranging from stringent on the left to permissive on the right. The response binary variable represents the odds of maintained or gained promoter interactions. The independent variables reflect the enrichment scores for 62 targets (TFs, histones and histone modifications), as well as the CHiCAGO score, the promoter interaction distance (log_10_ bp) and a boolean variable that indicates whether an interaction is between two baited promoters. The y-axis shows the regression beta which is positive for maintained promoter interactions and negative for lost promoter interactions. Vertical dotted line shows The value λ (λ1SE) at which the independent variables were tested for significance. The table on the right shows the FDR values at λ*_1SE_* for the significant predictors.

**Figure S4.**
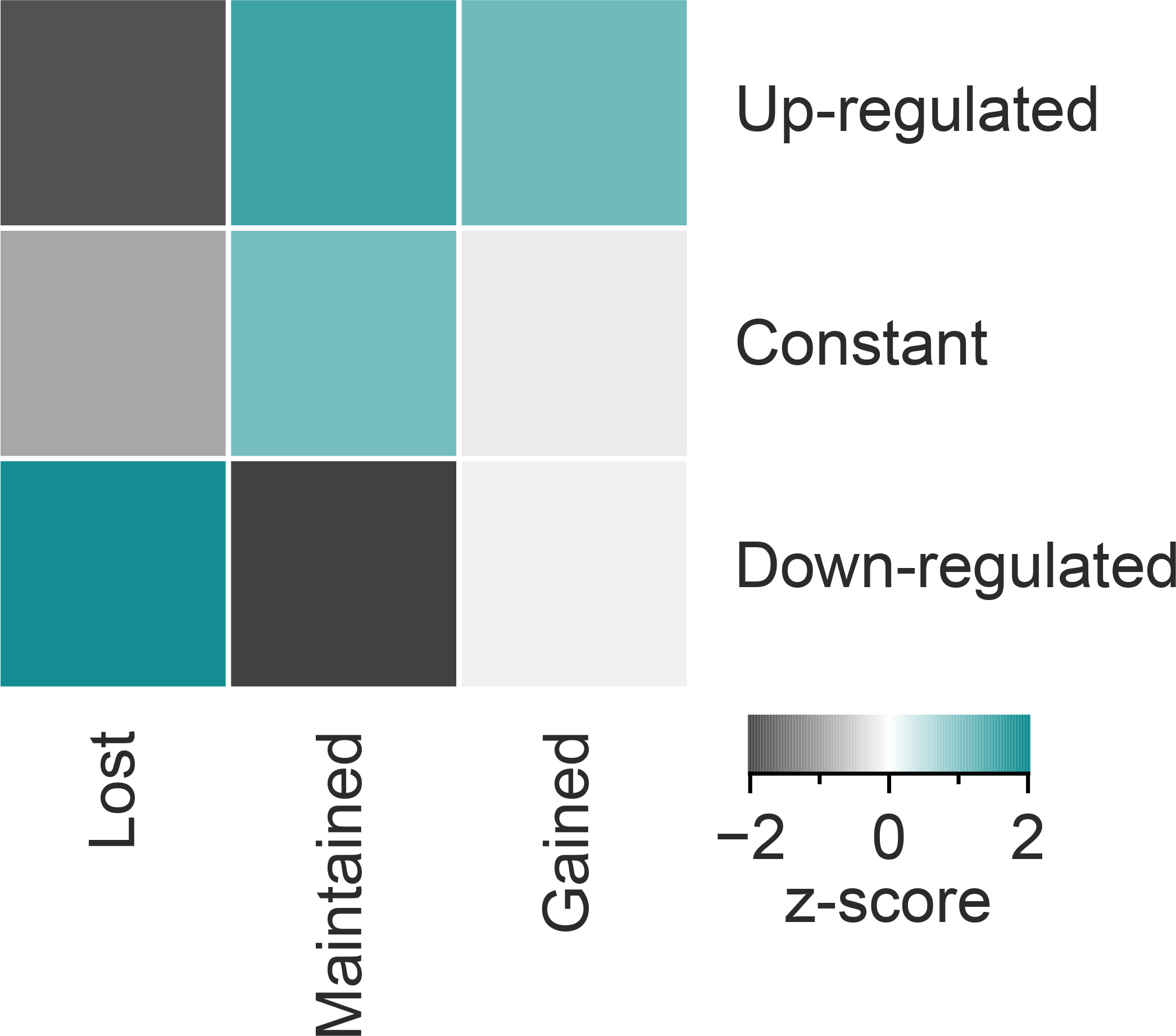
Association of transcriptional response upon SCC1 depletion associates with promoter interaction rewiring. Heatmap showing the z-scores of association between PIR interaction rewiring (columns) and transcriptional dynamics (rows) categories obtained from permutation analysis.

**Figure S5.**
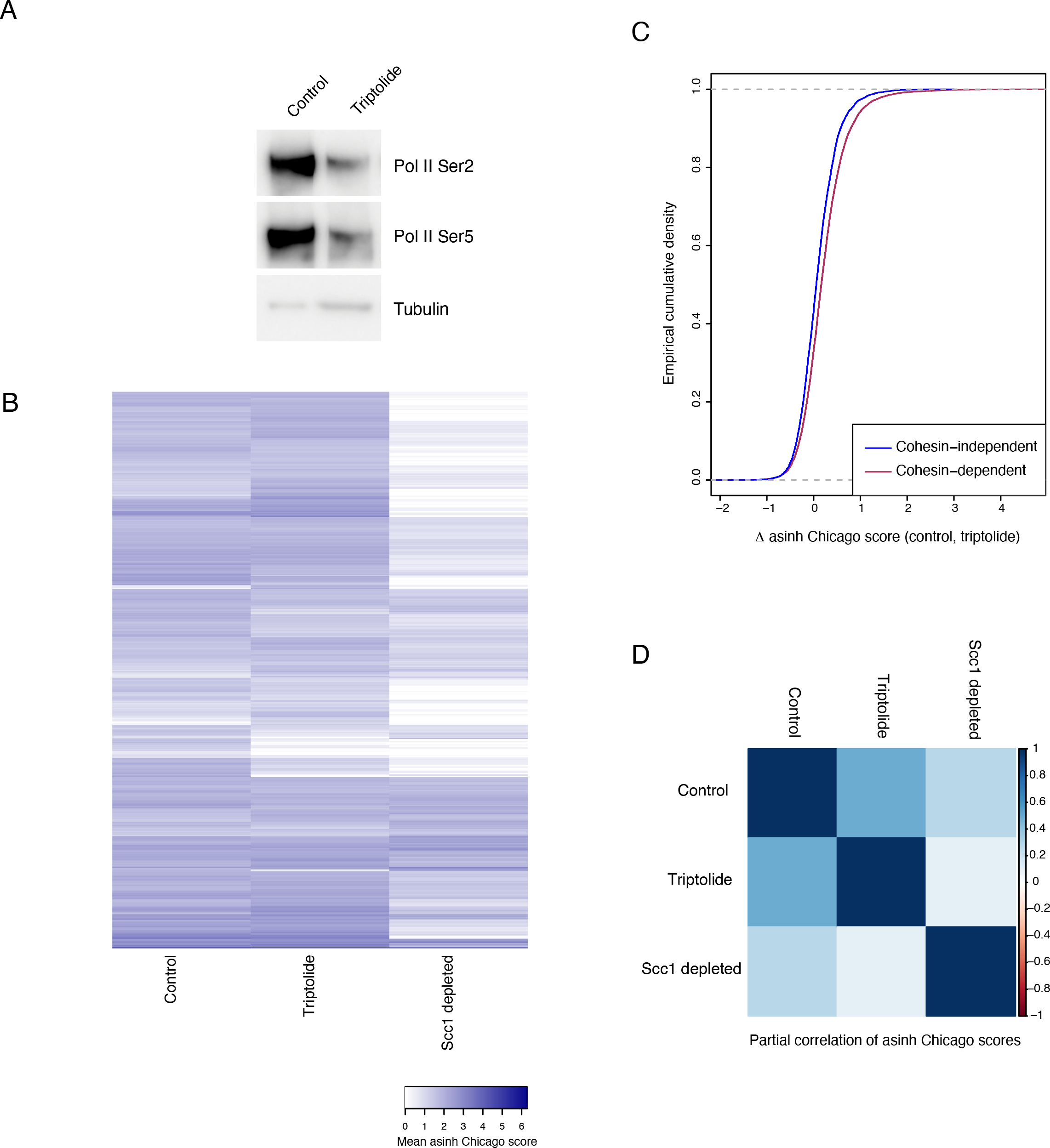
Triptolide treatment has a mild effect on promoter interactions. (A) Western blot analysis of control HeLa cells and those treated with 300nM triptolide for 3h using antibodies against RNA Pol II Ser2, RNA Pol II Ser5 and Tubulin as a loading control. (B) Heatmap showing the mean asinh-transformed Chicago score for 10,000 random promoter interactions detected in control HeLa cells, those treated with triptolide and SCC1-depleted cells. (C) Empirical cumulative density plot showing change in asinh-transformed Chicago scores upon triptolide treatment for cohesin-dependent (blue) and independent (red) interactions. Triptolide treatment has a relatively mild effect both interaction rewiring categories, with cohesin-independent interactions showing smaller-scale changes. (D) Heatmap showing partial correlations between asinh Chicago scores in the control, triptolide-treated and SCC1-depleted HeLa cells. Triptolide treated cells have positive partial correlations with both control cells and SCC1-depleted cells, with a much stronger effect for the former category. Jointly, these results do not favour the model of cohesin-independent interactions selectively stabilised by RNA Pol II activity or transcription.

